# Membrane-anchored mobile tethers modulate condensate wetting, localization, and migration

**DOI:** 10.1101/2024.12.04.626804

**Authors:** Qiwei Yu, Trevor GrandPre, Andrew G.T. Pyo, Andrej Košmrlj, Ned S. Wingreen

**Affiliations:** Lewis-Sigler Institute for Integrative Genomics, Princeton University, Princeton, NJ 08544; Department of Physics, Princeton University, Princeton, NJ 08544; Department of Physics, Washington University in St. Louis, MO, 63130; Department of Applied Physics, Stanford University, Stanford, CA 94305; Department of Mechanical and Aerospace Engineering, Princeton University, Princeton, NJ 08544; Princeton Materials Institute, Princeton University, Princeton, NJ 08544; Department of Molecular Biology, Princeton University, Princeton, NJ 08544

## Abstract

Biomolecular condensates frequently rely on membrane interactions for recruitment, localization, and biochemical substrates. Many of these interactions are mediated by membrane-anchored molecules such as proteins or specific lipids, which we refer to as “mobile tethers” since they can typically diffuse within the membrane while still interacting with the condensate. The presence of mobile tethers creates a surface with dynamic and spatially inhomogeneous wetting properties that are typically overlooked by traditional wetting theories. Here, we propose a general theoretical framework to study how mobile tethers impact both equilibrium and dynamic properties of condensate wetting. We show that a favorable tether-condensate interaction leads to tether enrichment at the condensate-membrane interface, which modifies the equilibrium surface tension and contact angle. Increasing tether abundance on the membrane can drive transitions between wetting regimes, with only a modest tether density and binding energy required for biologically relevant scenarios. Furthermore, tethers modulate how condensates react to complex membrane geometries. By helping condensates coat membranes, mobile tethers can facilitate condensate localization to junctions of membrane structures, such as the reticulated membranes inside the algal pyrenoid. Both tether abundance and mobility affect how droplets interact with complex membrane geometries, such as droplet migration along membrane tubules of varying radii. These results provide a framework to study the implications of tether-mediated condensate-membrane interactions for cellular organization and function.

## I. INTRODUCTION

Biomolecular condensates—intracellular compartments formed via phase separation—are essential for diverse biological processes, including gene regulation, metabolism, and cell signaling [1, 2]. In many instances, proper condensate function relies on interactions with membranes [3–8]. These membrane interactions can spatially organize condensates, concentrate interaction partners, and facilitate access to reactants. The algal pyrenoid exemplifies this interplay [9]: condensates enriched with the CO_2_-fixing enzyme Rubisco form around traversing membranes that supply CO_2_ to enhance photosynthetic efficiency. Conversely, condensates can also facilitate membrane processes such as transport, signaling, force generation, and structural remodeling. For example, Focal Adhesion Kinase (FAK) forms condensates on the cytoplasmic membrane, binding to lipids at sites where focal adhesions assemble, thereby regulating cell motility [10]. Similarly, B cell activation involves condensation on the plasma membrane that is essential for downstream signaling [11]. More broadly, unraveling the dynamic relationship between condensates and membranes is proving to be essential for understanding intracellular organization and function.

In many cases, membrane-associated condensates do not directly wet membranes. Instead, they adhere to membrane surfaces via tethering molecules, such as proteins or specific lipids, that are anchored to the membrane. In the pyrenoid of the model alga *Chlamydomonas reinhardtii*, for example, pyrenoid-traversing membranes feature tethers like RBMP1, RBMP2, and SAGA1, which directly bind to Rubisco [12, 13]. These tether proteins are essential for the assembly of the pyrenoid condensate around traversing membrane tubules, a structure that is crucial for the pyrenoid’s function in CO_2_ fixation. In this case and others, elucidating how tethers mediate condensate-membrane interactions is key to understanding the structure and function of membrane-associated condensates.

A key characteristic of these tether molecules is their ability to diffuse laterally within the membrane. As the condensate wets the membrane, the tethers can dynamically redistribute, enriching at the condensate-membrane interface due to favorable interactions with the condensate. This creates a surface with dynamic and spatially inhomogeneous wetting properties, which can affect both the equilibrium and dynamic aspects of condensate wetting. These effects are typically overlooked by traditional wetting theories, which often assume static surface properties [14, 15], or theories of soft wetting, where the dynamics comes from substrate deformation [16]. Here, motivated by both biological significance and theoretical interest, we seek to address the general question of how mobile tethers affect the condensate-membrane interaction and wetting.

In this work, we present a general theoretical framework that describes the coupled dynamics of condensates and mobile tethers. We find that mobile tethers enrich in the condensate-membrane interface, thereby reducing the surface tension with the membrane and modifying the equilibrium contact angle. By tuning the expression level of attractive tethers, cells can drive transitions from nowetting to partial or complete wetting. The per-tether binding energy required for such wetting transitions is estimated to be modest (only a few *k*_B_*T* ) for typical values of tether density and condensate surface tension. Furthermore, mobile tethers facilitate condensate local-V ization to intersecting membrane structures, such as the reticulated membranes inside the pyrenoid. Finally, both tether abundance and mobility affect droplet migration on spatially varying membrane structures such as tapering tubules. Overall, our framework provides tools for understanding the role of tether-mediated condensate-membrane interactions in cellular organization and function.

## II. RESULTS

### A. A general theoretical framework for tether-mediated wetting

We study a general theory that describes the densities of tethers and condensates with fields *ψ* and *ϕ*, respectively. A high (low) value of *ϕ* corresponds to a condensate dense (dilute) phase. The interactions are captured by a total free energy

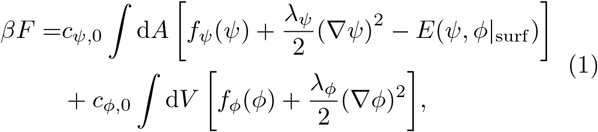

where the first integral is over the membrane area, and the second integral is over the bulk volume. Energy is measured in units of *β*^−1^ = *k*_B_*T*. *c*_*ψ*,0_ and *c*_*ϕ*,0_ are reference concentrations for the tether and condensate so that the free-energy densities are nondimensionalized: *E*(*ψ, ϕ* |_surf_ ) captures both condensatetether and condensate-membrane interactions; *f*_*ψ*_(*ψ*) and *f*_*ϕ*_(*ϕ*) are the free-energy densities of tethers and condensates respectively; *λ*_*ψ*_ and *λ*_*ϕ*_ are constants associated with interface energies.

The model encompasses a large class of systems and interactions by allowing the free-energy densities *f*_*ψ*_(*ψ*), *f*_*ϕ*_(*ϕ*), and the interaction energy *E*(*ψ, ϕ* |_surf_ ) to take any reasonable form. By minimizing the free energy in Eq. 1, we obtain the equilibrium concentration profile, from which the contact angle *θ* can be measured (Fig. 1A– B). To study the dynamics of wetting, we can further prescribe conserved (model B) dynamics [17]:

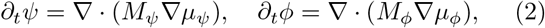

where *M*_*ψ*_ and *M*_*ϕ*_ are mobility coefficients, and *µ*_*ψ*_ = *δF/δψ* and *µ*_*ϕ*_ = *δF/δϕ* are the chemical potentials of the tethers and condensate, respectively.

**FIG. 1.**
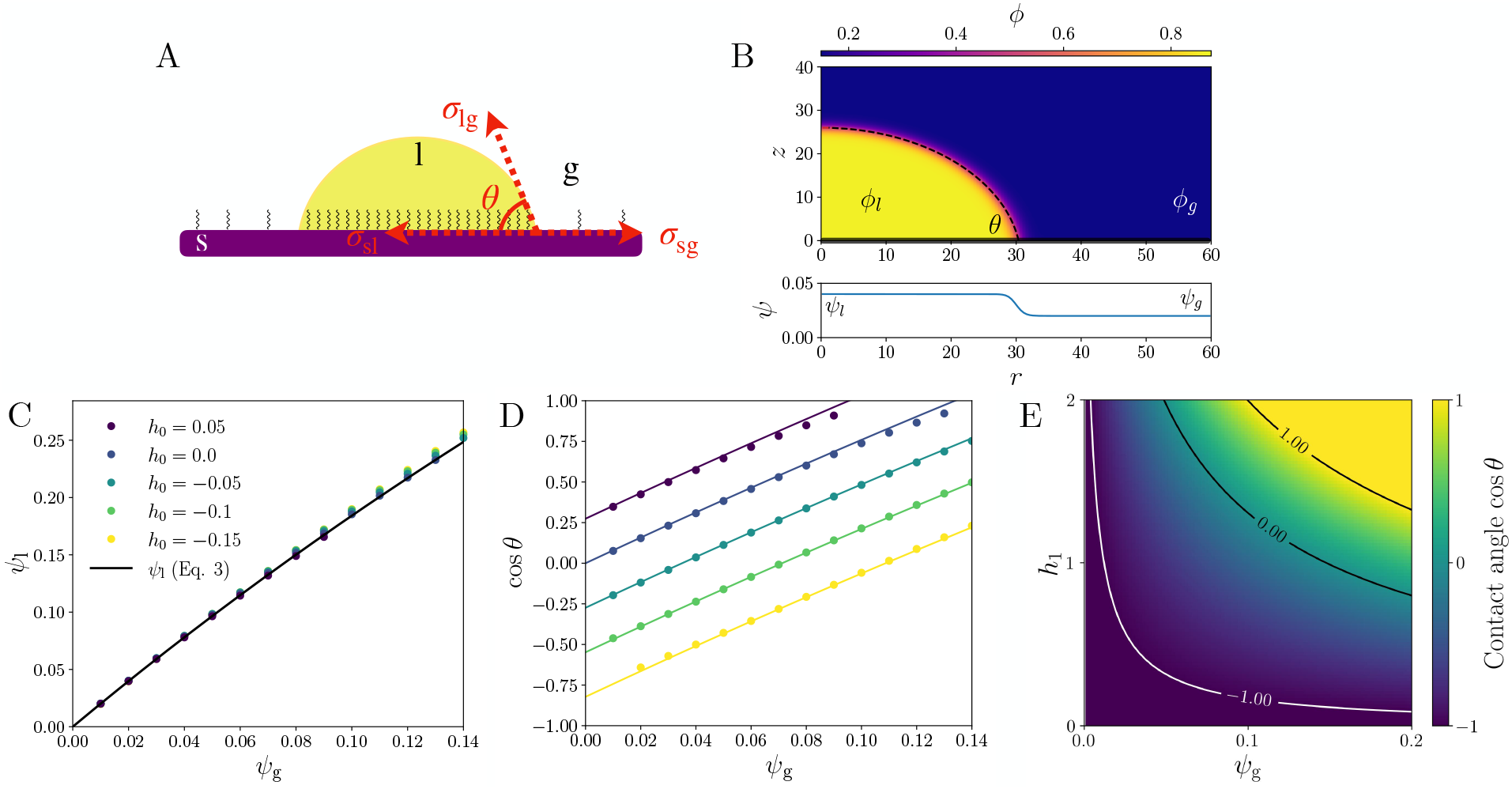
Mobile-tether-mediated condensate wetting of membranes. (A) Illustration of a biomolecular condensate (yellow) interacting with mobile tether molecules (black) to wet a membrane (purple). The interaction creates a localized enrichment of tethers around the condensate, surrounded by a lower background concentration of tethers. s, l, and g represent membrane (“solid”), dense phase (“liquid”), and dilute phase (“gas”). The contact angle *θ* is determined by force balance at the three-phase junction: *σ*_lg_ cos *θ* = *σ*_sg_ − *σ*_sl_, where the *σ*s are surface tensions. (B) A typical equilibrium concentration profile obtained from numerical simulations. The condensate field *ϕ* (top) and tether field *ψ* (bottom) are plotted in cylindrical coordinates (*r, z*) with axial symmetry. The thick black line indicates the flat membrane at *z* = 0. The black dashed curve is a spherical cap fit to the condensate surface contour. (C) Condensate-enriched tether concentration *ψ*_l_ increases with bulk tether concentration *ψ*_g_, for different condensate-membrane interactions *h*_0_, consistent with theory (solid curve, Eq. 3). (D) Contact angle cos *θ* as a function of tether concentration *ψ*_g_ for different *h*_0_ (see legend in C) agrees well with theory (solid curves, Eq. 4). (E) cos *θ* (Eq. 4) as a function of condensate-tether interaction *h*_1_ and tether concentration *ψ*_g_ for *h*_0_ = −0.2. Contours for cos = ±1 represent wetting transitions to complete and no wetting, respectively. In all simulations, *ψ* has a Dirichlet boundary condition while *ϕ* has a no-flux boundary condition. See *SI Appendix* for details and parameters.

To illustrate the physical picture, we study a minimal scenario of tether-mediated wetting. We consider a linear interaction energy *E*(*ψ, ϕ*) = (*h*_0_ + *h*_1_*ψ*)*ϕ*, where *h*_0_ and *h*_1_ describe condensate-membrane and condensate-tether interactions, respectively. We use Flory-Huggins free energies for self-energies *f*_*ξ*_(*ξ*) = *ξ* ln *ξ* + (1 −*ξ*) ln(1 −*ξ*) + *χ*_*ξ*_*ξ*(1 −*ξ*), with *ξ* ∈ {*ψ, ϕ*} representing the area or volume fraction of tether or condensate, respectively [18]. We set the units of free-energy densities via *c*_*ψ*,0_*k*_B_*T* = 1 and *c*_*ϕ*,0_*k*_B_*T* = 1, unit of length by *λ*_*ϕ*_ = 1, and unit of time by *M*_*ϕ*_ = 1. Further assuming non-self-interacting mobile tethers (*χ*_*ψ*_ = 0, *λ*_*ψ*_ = 0), we arrive at a minimal model for interrogating how tethers affect condensate wetting. We emphasize that the reported qualitative behaviors are generic and not sensitive to the specific choice of the functions for free-energy densities and condensate-tether interaction energy. It is also straightforward to extend the model to describe multi-component condensates and/or tethers, as well as more complex interactions.

### B. Mobile tethers control equilibrium wetting properties

In classical wetting theory, the contact angle *θ* of a droplet on a surface is determined by force balance at the three-phase junction through the Young-Dupré equation [14], which relates *θ* to the difference of surface tensions (Fig. 1A). In the presence of mobile tethers, however, favorable tether-condensate interactions enrich tethers within a wetting condensate (Fig. 1A), thereby creating a surface with inhomogeneous wetting properties, which in turn modifies the surface tensions and the contact angle.

Condensate phase separation creates dense and dilute phases in the bulk, with binodal concentrations *ϕ*_l_ and *ϕ*_g_ (as measured in volume fractions), respectively. The concentration difference Δ*ϕ* = *ϕ*_l_ − *ϕ*_g_ drives the attraction of tethers to the condensate, resulting in a tether area fraction *ψ*_l_ in contact with the dense phase, which is higher than that in contact with the dilute phase *ψ*_g_ (Fig. 1B). This partition of tethers reaches equilibrium when chemical potentials are balanced: *µ*_*ψ*,l_ = *µ*_*ψ*,g_, where *µ*_*ψ*,∗_ = *δF/δψ*_∗_ for ∗ ∈ {l, g}, which leads to (see *SI Appendix* for details)

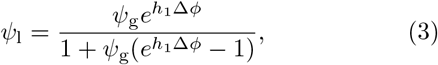

where we have approximated the condensate concentrations at the surface with the bulk binodal concentrations.

This agrees well with numerical simulations across a wide range of *ψ*_g_, for both repelling (*h*_0_ *<* 0) or attracting (*h*_0_ *>* 0) interactions between the bare membrane and the condensate (Fig. 1C).

The presence of attractive tethers reduces both surface tensions *σ*_sl_ = ln(1 −*ψ*_l_) −*h*_0_*ϕ*_l_ and *σ*_sg_ = ln(1 −*ψ*_g_) − *h*_0_*ϕ*_g_. However, the decrease in *σ*_sl_ is more substantial due to tether enrichment in the condensate (*ψ*_l_ *> ψ*_g_). This, in turn, modifies the contact angle *θ*, which is determined by force balance at the three-phase junction: *σ*_lg_ cos *θ* = *σ*_sg_ −*σ*_sl_. The modified contact angle is (see *SI Appendix* for details)

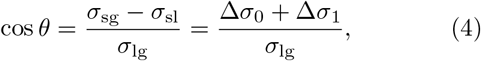

where Δ*σ*_0_ = *h*_0_Δ*ϕ* is the surface tension difference in the absence of tethers, and 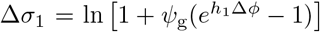 is the additional surface tension difference due to mobile tethers. Δ*σ*_1_ increases monotonically with tether abundance *ψ*_g_ and tether-condensate interaction *h*_1_. Indeed, numerical simulations find the contact angle in simulations to be in excellent agreement with Eq. 4 (Fig. 1D, solid curves). Thus, an attractive interaction due to mobile tethers can substantially modulate wetting over a wide range of contact angles.

Wetting transitions occur at cos *θ* = 1, when a droplet completely wets a membrane, and at cos *θ* = −1, when a droplet detaches from a membrane (non-wetting). Tethers can induce transitions between these wetting regimes: For a repelling membrane that is initially in the non-wetting regime (*h*_0_ *<* − *σ*_lg_*/*Δ*ϕ*), both partial wetting [cos *θ*∈ ( − 1, 1)] and complete wetting (cos *θ* = 1) regimes can be achieved via a high enough density of attractive tethers (Fig. 1E). To reach complete wetting, the required critical density of tethers is 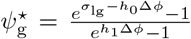, which must stay below 1 since *ψ* is defined in terms of area fraction. Since 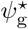 vanishes in the limit of large *h*_1_, a finite density of tethers is sufficient to access all three wetting regimes as long as the tether-condensate attraction is strong enough.

For real tether molecules, how much binding energy is required to significantly affect wetting properties? Typically, the membrane would be slightly repulsive for polymer condensates because being close to a membrane reduces the conformational entropy of polymers, leading to an estimated Δ*σ*_0_ ∼ − 10^−1^*k*_B_*T/*nm^2^ [19]. In aqueous buffer, biomolecular condensate surface tensions are typically of the same order *σ*_lg_ ∼ 10^−1^*k*_B_*T/*nm^2^ [20]. Thus, to drive wetting, tethers must reduce surface tension by the same order Δ*σ*_1_ ∼ 10^−1^*k*_B_*T/*nm^2^. A typical tether density of *n*_g_ ∼ 10^−2^nm^−2^ [21] yields a required binding energy of *ϵ* ≈ 𝒪 (1)*k*_B_*T* (see *SI Appendix* for details).

Despite being a rough estimate, these calculations show that a modest per-tether binding energy (a few *k*_B_*T* ) is sufficient to drive wetting transitions. Therefore, cells can potentially regulate condensate wetting by tuning the expression level of tether molecules.

### C. Mobile tethers facilitate condensate localization dynamics

Thus far, we have focused on equilibrium morphologies. How might mobile tethers affect the dynamics of condensate formation and localization? In the alga *C. reinhardtii*, for example, the pyrenoid condensate dissolves and reforms every cell division [22], and the new pyrenoid centers around a reticulated region where many membrane tubules meet. Since the reticulated region has a high membrane area per volume, it might therefore be able to enrich tethers more effectively than other regions of the tubule. Hence, we hypothesize that mobile tethers may facilitate condensate localization by enrichment in the reticulated region.

To simply illustrate this mechanism, we study a two-dimensional system which is bounded by membranes on the left and bottom sides and closed on the other two (Fig. 2). The bottom-left corner is most favorable for the condensate since it can interact there with the largest amount of membrane area (and therefore tethers), analogous to the reticulated region in the pyrenoid. Initially in simulations, the condensate coats part of the membrane, and its bulk concentration is between binodal and spinodal concentrations. If tethers have a high mobility, they quickly enrich in the condensate and help it localize to the corner (Fig. 2A). In contrast, if the tether mobility is low, the condensate first breaks up into smaller droplets, and only slowly relocalizes to the corner through a coarsening process (Fig. 2B). Even though both reach the same equilibrium state, the latter process is much slower (Fig. 2C). Thus, by helping the condensate to optimize its membrane contacts, mobile tethers can facilitate coarsening and localization with respect to membrane structures.

**FIG. 2.**
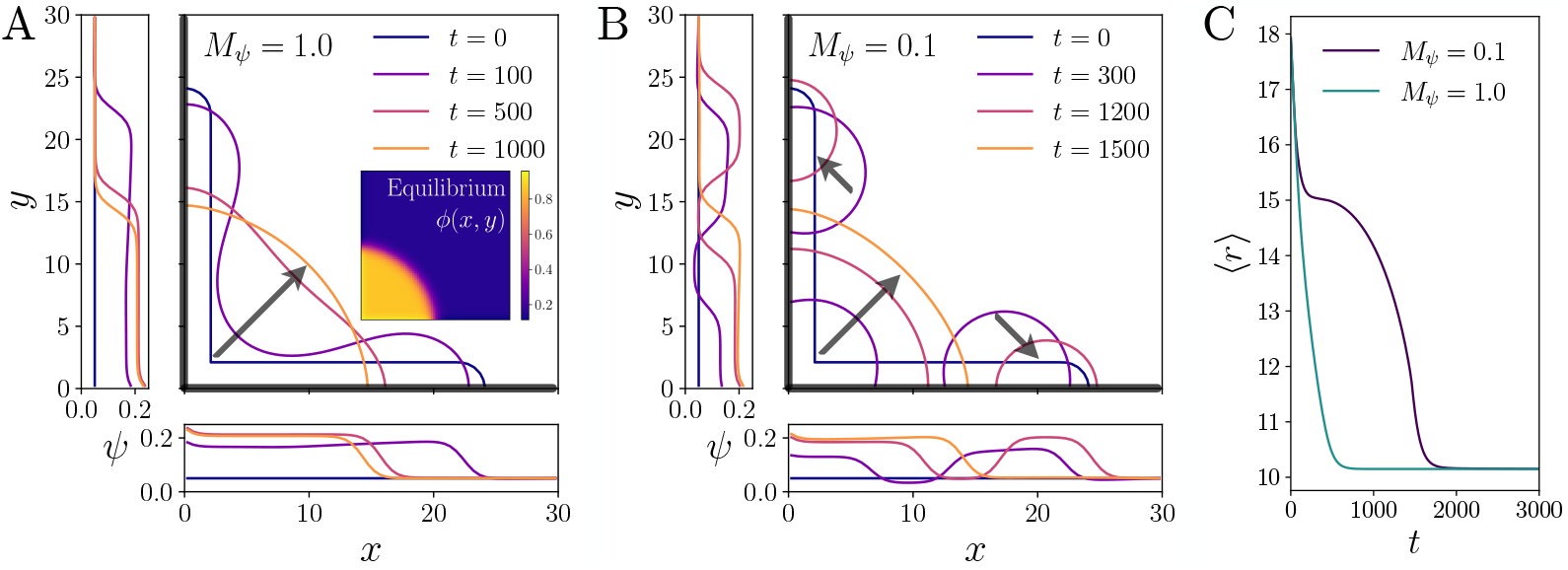
Mobile tethers facilitate dynamic condensate relocalization. (A–B) Dynamics of condensate localization for tether mobility *M*_*ψ*_ = 1.0 (A) and *M*_*ψ*_ = 0.1 (B). The simulation domain is a 2D system (*x, y*) with membranes on the left and bottom boundaries (indicated by thick black lines). Different colors indicate concentration profiles at different times (legend), with the condensate *ϕ* represented by interface contours and the tether density *ψ* shown in the left and bottom insets. Inset in (A) shows the final equilibrium profile for *ϕ*(*x, y*). Black arrows indicate the time evolution of the interface contours to guide the eye. The tether density at the boundaries is *ψ*_g_ = 0.05. The overall ⟨*ϕ*⟩ is conserved due to no-flux boundary conditions. (C) Condensate location as quantified by the average distance from the bottom-left corner location as quantified by the average distance from the bottom-left corner 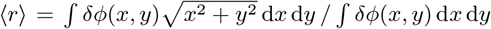 where *δϕ* = *ϕ* − *ϕ*_g_. See *SI Appendix* for details and simulation videos.

### D. Tether abundance and mobility affect condensate migration on tubules

Our theoretical framework enables the study of mobiletether-mediated wetting of a myriad of possible membrane structures, including tubes, sheets, and cristae. As highlighted in the example above, the presence of mobile tethers could modulate or amplify the effects of membrane geometry on condensate behavior.

To illustrate such geometric effects, we consider the dynamics of a condensate that wets a membrane tubule of varying radius. Here we consider a (truncated) cone geometry where the tubule radius varies linearly along its long axis (Fig. 3A, black line), although the theoretical arguments are general for other geometries as well. When the tubule is thin (compared to *V* ^1/3^, where *V* is the droplet volume), the droplet can adopt an axisymmetric barrel-like shape that wraps around the tubule. By contrast, the droplet can also wet only one side of the tubule and adopt an asymmetric clamshell-like shape

**FIG. 3.**
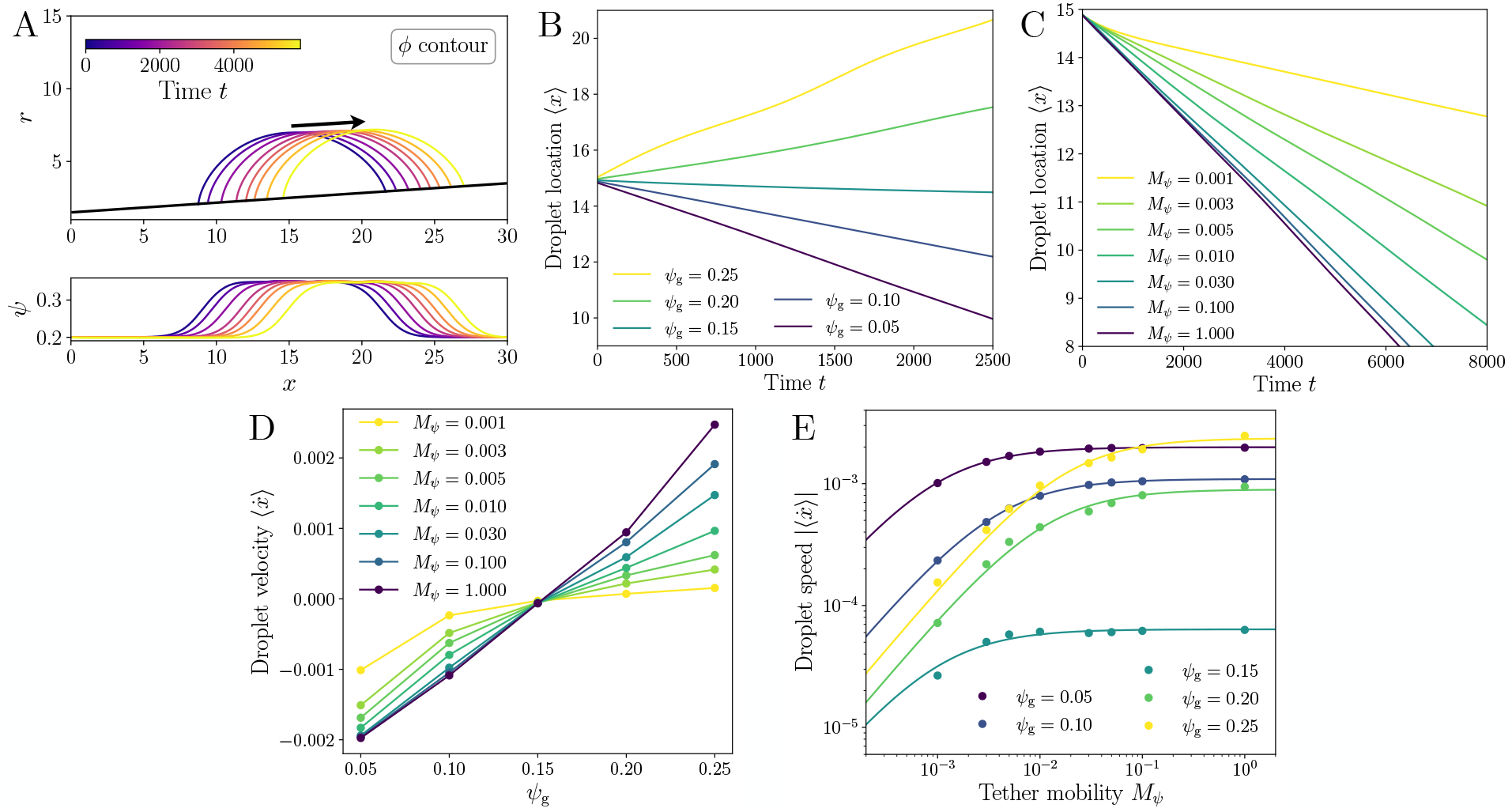
Mobile tethers affect condensate migration on a tubule of varying radius. Here, we consider a (truncated) cone geometry where the tubule radius varies linearly along its long axis, although similar results hold for other geometries as well. (A) Time course of the condensate *ϕ* (top contours) and tether *ψ* (bottom) densities on a tubule of varying radius, in cylindrical coordinates (*r, x*) where *x* runs along the central axis of the tubule. Curves of different colors represent different times (inset legend) with the arrow indicating the direction of migration. The black line indicates the tubule surface. (B–C) Condensate location as quantified by the average position ⟨*x*⟩ for different tether concentrations *ψ*_g_ for *M*_*ψ*_ = 1.0 (B), and different tether mobilities *M*_*ψ*_ for *ψ*_g_ = 0.10 (C). (D) Migration velocity 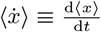as a function of *ψ*_g_ for different *M*_*ψ*_. (E) Migration speed 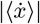 as a function of *M*_*ψ*_ for different *ψ*_g_. Solid curves are fits to 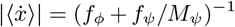 with *f*_*ϕ*_ and *f*_*ψ*_ as fitting parameters. See *SI Appendix* for details and parameters.

when the tubule is thick [23–26]. Here, we focus on the former case, where the droplet is able to wrap around the tubule (Fig. 3A).

We expect such an axisymmetric droplet to migrate along the tubule, moving down the gradient of free energy until reaching a minimum-energy equilibrium position. The equilibrium location will depend on the contact angle *θ*, where a smaller *θ* (more wetting) favors regions of larger radius, and vice versa. By approximating the cross-section of the barrel-shaped droplet as circular (see *SI Appendix* for numerical justification), we find that the droplet always moves to the smallest radius for *θ > π/*2, while for *θ < π/*2 the droplet prefers a finite radius that scales as *r*∼ *V* ^1/3^ cot *θ* (see *SI Appendix* for details). We note, however, that if *r/V* ^1/3^ is too large, the axisymmetric barrel becomes unstable and the droplet moves to wet only one side of the cylinder (clamshell shape) [23, 26]. Nevertheless, for a droplet initialized on a relatively thin tubule, the contact angle *θ* dictates whether it initially moves to small or large radius.

Since the contact angle *θ* can be modulated by tether abundance *ψ*_g_ (Eq. 4, Fig. 1D), we expect that *ψ*_g_ can affect the equilibrium location of the droplet on the tubule. Specifically, increasing tether abundance *ψ*_g_ decreases *θ* (Fig. 1D), thereby shifting the equilibrium location to a larger radius. Indeed, when we initialize a droplet at a particular location on the tubule, it migrates towards small radius when *ψ*_g_ is low (large *θ*), but towards large radius when *ψ*_g_ is high (small *θ*) (Fig. 3B). Increasing tether mobility *M*_*ψ*_ leads to faster migration (Fig. 3C), while a very small *M*_*ψ*_ can lead to self-trapping, pinning the droplet and arresting migration.

These results suggest that tether abundance and mobility affect different aspects of droplet migration on spatially varying membrane structures: Tuning tether abundance *ψ*_g_ can modulate the total force on the droplet and control its preferred localization on the tubule, while tuning tether mobility *M*_*ψ*_ can control droplet migration speed (Fig. 3D). In the overdamped limit, the driving force due to the free-energy gradient (or equivalently, surface tension forces) is balanced by viscous drag from both the condensate and the tethers. Here, the drag is controlled by the mobility coefficients *M*_*ϕ*_ and *M*_*ψ*_. Thus, the droplet velocity is given by:

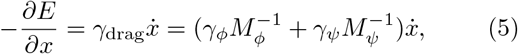

where the driving force 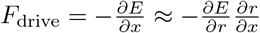 is due to the gradient of the energy of a droplet wetting a tubule of varying radius *r* (see *SI Appendix* for details); 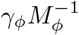and 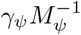 represent the drag due to the condensate and the tethers, respectively. Thus, droplet speed depends on tether mobility via an inverse linear relationship 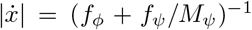, with 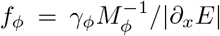and *f*_*ψ*_ = *γ*_*ψ*_*/* |*∂*_x_*E* |. The ratio of coefficients *f*_*ψ*_*/f*_*ϕ*_ = *M*_*ϕ*_*γ*_*ψ*_*/γ*_*ϕ*_ depends on tether concentrations *ψ*_g_ and *ψ*_l_ (see *SI Appendix* ). This relation is in good agreement with numerical simulations (Fig. 3E). In other words, tethers can slow down droplet migration if they cannot redistribute quickly enough to maintain an energetically favorable wetting configuration as the droplet moves. In the limit of immobile tethers (*M*_*ψ*_ → 0), the droplet becomes trapped in place.

Taken together, our results show that mobile tethers provide a mechanism to control how condensates respond to membrane geometry by modulating both the condensate’s favorable location and its migration speed.

## III. DISCUSSION

Membrane proteins and specialized lipids play an important role in regulating membrane functions, including their interaction with biomolecular condensates. However, the mobility of tethering molecules within the membrane has been largely overlooked in previous studies of condensate wetting. Here, we develop a general theoretical framework for mobile-tether-mediated wetting and show that tethering molecules can substantially modulate both equilibrium and dynamical aspects of condensate wetting, including migration and localization. These results suggest potential mechanisms for cells to regulate condensate formation and organization via the expression of mobile tethering molecules.

Our theory is relevant for a wide range of biological systems, including the algal pyrenoid [12, 13, 27], focal adhesion proteins [28, 29], T-cell activation [30], actin assembly [31], and potentially the organization of ER exit sites [32–34]. Since a modest tether density and per-tether binding energy (a few *k*_B_*T* ) would be sufficient to substantially affect wetting properties, it is plausible for cells to regulate a wide range of condensates via mobile tethering molecules. Experimentally perturbing tethering molecules in cells will provide valuable insights into their importance for these structures.

Besides providing insights into *in vivo* structures and functions, our framework also makes quantitative predictions that can be tested *in vitro*. One direct test would be to place fluorescently tagged tethering molecules in supported lipid bilayers (SLBs) and track the tether concentrations *ψ*_g_ and *ψ*_l_ as the membrane is wetted by a condensate that is attracted to the tether molecule. Repeating such experiments at different tether concentrations *ψ*_g_ would enable a quantitative test for the tether enrichment predicted by theory (Eq. 3 and Fig. 1C). In addition, the contact angle can potentially be measured by confocal imaging and compared with theory (Eq. 4 and Fig. 1D).

While this work focuses on tether-mediated wetting of a fixed membrane, our framework can be extended to include effects such as membrane deformation [35, 36] and hydrodynamic coupling, as well as active processes [37], such as post-translational modification upon wetting. In a biological context, it will also be interesting to study how tether-mediated wetting affects downstream signaling, which is often a nonequilibrium process [38, 39]. Overall, our framework paves the way for the study of how mobile-tether-mediated interactions affect condensate morphology, dynamics, and function.

## ACKNOWLEDGMENTS

We thank Benny Davidovitch, Martin Jonikas, Alejandro Martínez-Calvo, Narayanan Menon, Pedro de Souza, and Hongbo Zhao for useful discussions. This work was supported by NSF grant MCB-2410354, by the Center for the Physics of Biological Function (NSF PHY-1734030), and by the Princeton Center for Complex Materials (NSF DMR-2011750). The work by Q.Y. was supported in part by a Harold W. Dodds Fellowship. The work by A.G.T.P was supported in part by a Stanford Science Fellowship. This research was supported by the National Institutes of Health under award number R01GM140032. The content is solely the responsibility of the authors and does not necessarily represent the official views of the National Institutes of Health.

## Supplemental Material

### I. THEORETICAL FRAMEWORK

As discussed in the main text, we consider a system where (three-dimensional) biomolecular condensates can interact with a (two-dimensional) membrane. The total free energy reads:

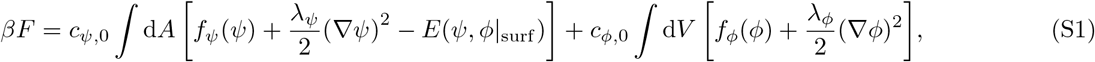

where *ϕ* is the condensate density field and *ψ* is the tether density field. The first integral is over the membrane area, while the second integral is over the bulk volume. *f*_*ϕ*_(*ϕ*) and *f*_*ψ*_(*ψ*) are the free-energy densities of the condensate and tethers, respectively; *E*(*ψ, ϕ* _surf_ ) describes the interaction energy between the condensate and the tether/membrane, with *ϕ* |_surf_ denoting the condensate density at the membrane surface. *λ*_*ψ*_ and *λ*_*ϕ*_ are related to the line/surface tensions. The free energy is measured in units of *β*^−1^ = *k*_B_*T*. The free-energy densities (integrands in Eq. S1) are non-dimensionalized by the factor of *β* and reference concentrations *c*_*ψ*,0_ and *c*_*ϕ*,0_, for the tether and the condensate, respectively [1]. For qualitative analysis and for the sake of simplicity, we set *c*_*ψ*,0_*k*_B_*T* = 1 and *c*_*ϕ*,0_*k*_B_*T* = 1; this does not affect the results but must be revisited when estimating the energy scales for real tethers (Sec. II).

To minimize the free energy, we prescribe the following gradient (model-B) dynamics for both the tethers and condensate [2]:

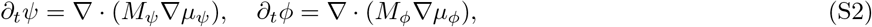

where *M*_*ψ*_ and *M*_*ϕ*_ are mobility coefficients, while *µ*_*ψ*_ = *δF/δψ* and *µ*_*ϕ*_ = *δF/δϕ* are the chemical potentials of the tethers and condensate, respectively. The condensate obeys no-flux boundary condition at the membrane surface:

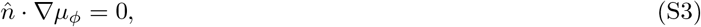

where 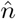is the unit normal vector pointing out of the membrane. Additionally, the bulk-surface interaction gives the following wetting boundary condition:

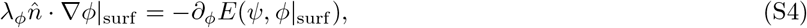

which reflects a local change in condensate concentration near the membrane surface due to the condensate’s interaction with the membrane and tethers.

While the model can be used to describe a large class of systems by allowing the free-energy densities *f*_*ϕ*_(*ϕ*) and *f*_*ψ*_(*ψ*) and the interaction energy *E*(*ψ, ϕ* |_surf_ ) to take different forms, we will focus on a minimal model to illustrate the essential physical picture.

#### A. A minimal model for tether-mediated wetting

For the sake of simplicity, we use Flory-Huggins free-energy densities for both the condensate and the tethers

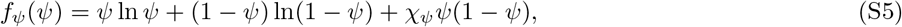

**FIG. S1.**
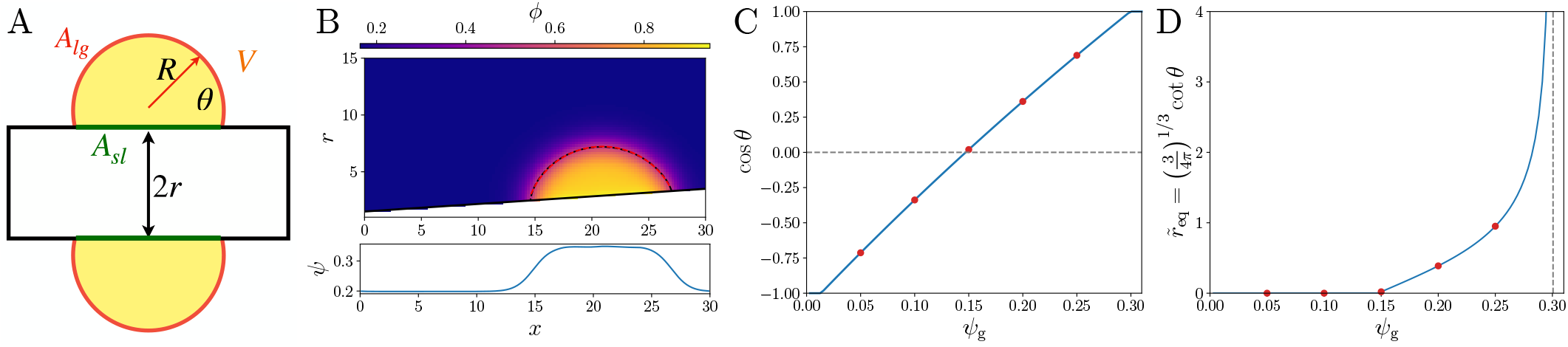
Equilibrium position of a droplet on a tubule of varying radius. (A) Illustration of the ansatz used for computing the free energy of a droplet of volume *V* wetting a cylinder of radius *r* with contact angle *θ*. This is a cross-sectional view where the droplet assumes a circular cap shape with radius *R*. (B) Snapshot of the condensate profile *ϕ* (top) and tether profile *ψ* (bottom) for a droplet migrating on a tubule (membrane indicated by the black line) obtained from numerical simulation. The red dashed curve shows that a circular fit is in good agreement with the interface contour (black dashed curve). (C) Contact angle cos *θ* as a function of tether density *ψ*_g_. (D) Normalized equilibrium tubule radius 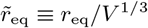as a function of tether density *ψ*_g_. In (C) and (D), blue curves are theoretical predictions (Eq. S26 and Eq. S38), and red data points simply indicate the theoretical predictions evaluated at the *ψ*_g_ values used in Fig. 3 in the main text. Parameters: *h*_0_ = − 0.2, *h*_1_ = 1, *χ*_*ϕ*_ = 2.5, *λ*_*ϕ*_ = 1, *χ*_*ψ*_ = 0, *λ*_*ψ*_ = 0. For (B), *ψ*_g_ = 0.20 and mobilities *M*_*ϕ*_ = 1.0 and *M*_*ψ*_ = 1.0.

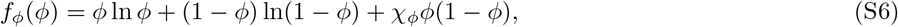

with *ψ* and *ϕ* denoting the area or volume fraction of tether and condensate, respectively. *χ*_*ψ*_ and *χ*_*ϕ*_ are the Flory-Huggins interaction parameters for the tether and the condensate, respectively.

We further assume a linear interaction energy (non-dimensionalized in the same way as *f*_*ψ*_)

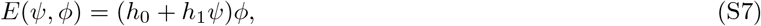

where *h*_0_ and *h*_1_ describe condensate-membrane and condensate-tether interactions, respectively. The chemical potentials are given by:

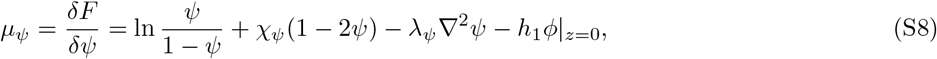

**FIG. S2.**
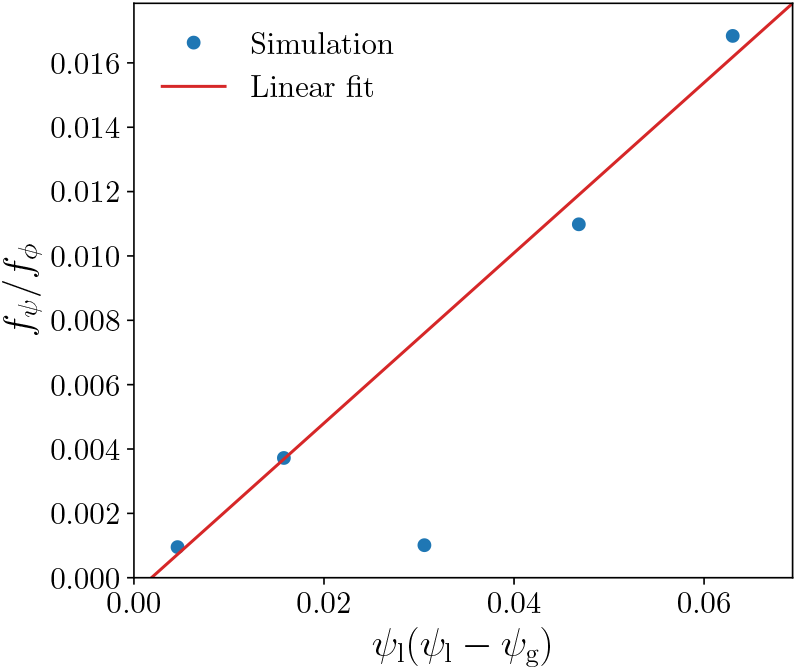
Proportionality between *f*_*ψ*_*/f*_*ϕ*_ and (*ψ*_l_ −*ψ*_g_)*ψ*_g_ (Eq. S52). Blue circles are numerical results obtained from simulations in Fig. 3E, and the red line is a linear fit excluding the point at *ψ*_g_ = 0.15, for which the droplet velocity is too small for an accurate estimate of the drag coefficients.

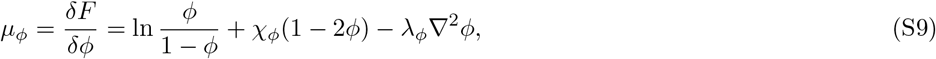

where *z* is the perpendicular distance from the membrane with *z* = 0 indicating the membrane surface. Further assuming for simplicity that tethers do not interact with each other and only interact with the condensate, we have *χ*_*ψ*_ = 0 and *λ*_*ψ*_ = 0. The tether chemical potential then simplifies to

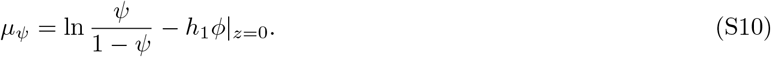

Let *ϕ*_l_ and *ϕ*_g_ denote condensate densities in the dense (liquid) phase and the dilute (gas) phase, respectively. The corresponding tether densities in the dense and dilute phases are denoted as *ψ*_l_ and *ψ*_g_. They are related to each other through chemical potential balance:

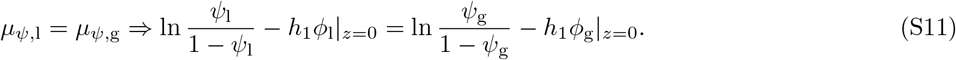

Approximating the surface density *ϕ*|_z=0_ with the bulk binodal values^1^, we have

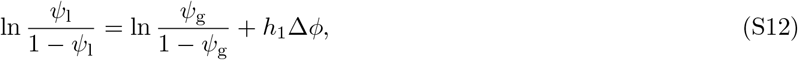

where Δ*ϕ* = *ϕ*_l_|_z=0_ − *ϕ*_g_|_z=0_ ≈ *ϕ*_l_ − *ϕ*_g_ is the difference between binodal concentrations. Solving for *ψ*_l_:

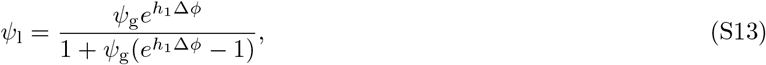

which is Eq. (3) in the main text.

The contact angle *θ* is given by force balance at the three-phase junction:

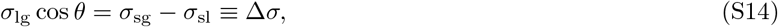

where the *σ*s represent surface tensions. *σ*_lg_ is the surface tension between the dense and dilute phases in 3D and is independent of the tether concentration. The surface tension with the membrane *σ*_s∗_, which depends on the tether density *ψ*, is given by computing the excess free energy (per unit area):

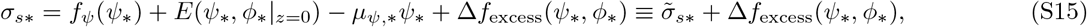

where ∗ ∈ {l, g} denotes the dense (“liquid”) or dilute (“gas”) phase, respectively. 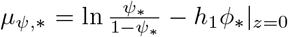 is the tether chemical potential. Δ*f*_excess_(*ψ*_∗_, *ϕ*_∗_) is defined below and will prove to be higher order in *h*_0_ and *h*_1_. We define 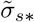 to be the sum of the first three terms, which is given by

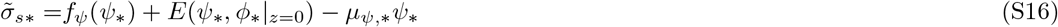

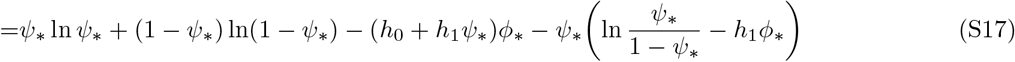

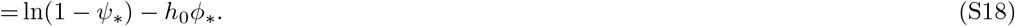

The surface tension difference due to these terms reads

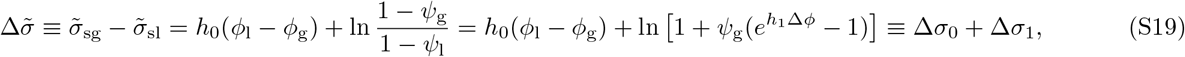

wh ere Δ*σ*_0_ = *h*_0_(*ϕ*_l_ − *ϕ*_g_) = *h*_0_Δ*ϕ* is the surface tension difference in the absence of tethers, and 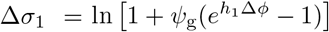 is the additional surface tension difference due to mobile tethers. We have substituted *ψ*_l_ with Eq. S13. Note that to the leading order, we have 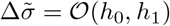.

Δ*f*_excess_(*ψ*_∗_, *ϕ*_∗_|_z=0_) is the excess free-energy density due to a boundary layer of condensate at the membrane surface. Let 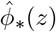denote the condensate concentration at distance *z* from the membrane in the dense (∗ = l) and dilute (∗ = g) phases. Near the membrane, 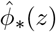 deviate from the binodal concentrations *ϕ*_∗_, leading to excess free energy per surface area:

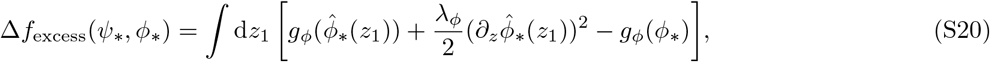

where *ϕ*_∗_ denotes the binodal concentration, and *g*_*ϕ*_(*ϕ*) = *f*_*ϕ*_(*ϕ*) − *µ*_*ϕ*_*ϕ* is the Gibbs free energy of the condensate. The concentration profile 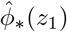 is the solution to the following boundary value problem:

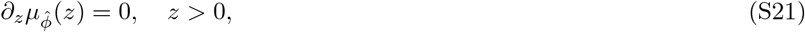

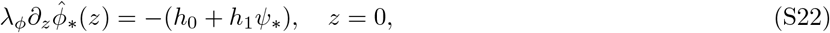

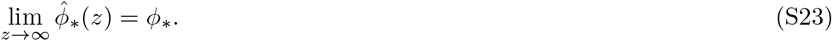

In practice, we find that the excess free energy density is negligible compared to the other terms. This can be explained by the following scaling argument: To the leading order in 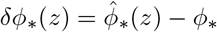, the excess free energy is

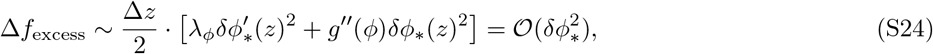

where Δ*z* is the boundary layer thickness, and we have used *g*^*′*^(*ϕ*) = 0 at the binodal concentration. From the wetting condition [Eq. S22], we find *δϕ*_∗_ = (*h*_0_, *h*_1_). Hence, the excess free energy becomes quadratic in the interaction parameters:

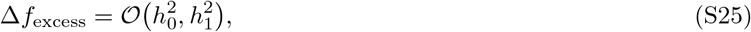

which is a higher-order term compared to 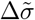.

Putting everything together:

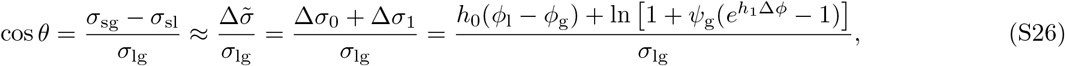

which produces Eq. (4) in the main text.

To achieve complete wetting (cos *θ* = 1), the critical tether density 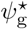 is given by

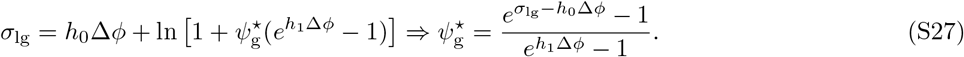

### II. ESTIMATING THE TETHER BINDING ENERGY REQUIRED TO DRIVE WETTING TRANSITION

Here, we estimate the value of Δ*σ*_0_, which is the dilute/dense phase surface tension difference due to the condensate-membrane interaction. Typically, a membrane would be slightly repulsive for a polymer condensate because being close to the membrane reduces the conformational entropy of the polymers. Thus, we can estimate the magnitude of this effect by considering polymer “blobs” close to the membrane. Each “blob” would contribute 1*k*_B_*T*, and the number of polymer “blobs” per unit area could be estimated by 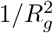with *R*_g_ being the radius of gyration. For an intrinsically disordered protein (IDP) of length ∼ 100 a.a., we estimate *R*_g_ ∼ 3nm [3], and therefore 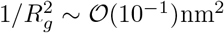. Thus, the surface tension due to entropic repulsion is of the order Δ*σ*_0_∼−𝒪 (10^−1^)*k*_B_*T/*nm^2^.

On the other hand, previous micropipette aspiration found that the typical biopolymer condensate surface tension *σ*_lg_ is at most 𝒪 (1)mN*/*m, or equivalently 𝒪 (10^−1^)*k*_B_*T/*nm^2^ [4]. Thus, to achieve complete wetting, the additional surface tension difference due to tethers must also reach Δ*σ*_1_ = *σ*_lg_ − Δ*σ*_0_∼𝒪 (10^−1^)*k*_B_*T/*nm^2^.

To estimate the corresponding binding energy relevant for real tethers, we recall that the surface tension was renormalized by *c*_*ψ*,0_*k*_B_*T*, with a close-packed tether density set by *c*_*ψ*,0_. Thus, in the limit of dilute tethers *ψ*_g_ = *c*_g_*/c*_*ψ*,0_ ≪ 1, where *c*_g_ is the (dimensional) tether density in contact with the dilute phase, we have

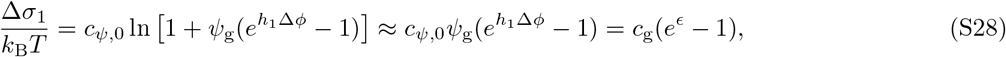

where *c*_g_ ∼10^−2^nm^2^ [5] is the dimensional tether number density, and *ϵ* = *h*_1_Δ*ϕ* is the energy reduction per tether when inside the condensate, measured in units of *k*_B_*T*. This leads to

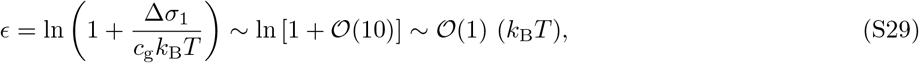

which suggests that a binding energy of a few *k*_B_*T* per tether is sufficient to modulate equilibrium condensate-membrane wetting properties.

### III. DROPLET MIGRATION ON A TUBULE OF VARYING RADIUS

#### A. Equilibrium position of a droplet

In this section, we consider the free energy of a droplet as it wets a membrane tubule in an axisymmetric configuration that wraps around the tubule. If the radius of the tubule is slow-varying compared to the droplet size, we can approximate the tubule locally as a cylinder of radius *r* (Fig. S1A). Thus, the free energy of the droplet *E*(*V, r*) is determined by its volume *V* and the local tubule radius *r*.

The droplet adopts a barrel-shaped constant-mean-curvature surface when wetting the cylinder [6]. For analytical tractability and motivated by our numerical simulations (Fig. S1B), we approximate the cross-section profile as a circular cap (Fig. S1A). The radius *R* is determined by constraining the volume of the droplet:

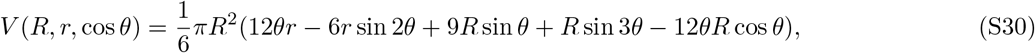

where *θ* is the contact angle. The total surface free energy is given by

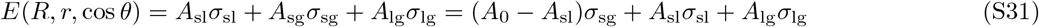

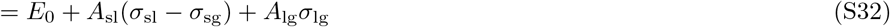

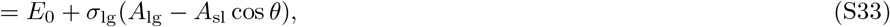

where *A*_0_ is the total membrane area, *A*_sl_, *A*_sg_, and *A*_lg_ are the areas of the membrane-condensate, membrane-solvent, and condensate-solvent interfaces, respectively. The last line follows from the force balance at the three-phase junction (Eq. S14). Applying our droplet shape ansatz, we have

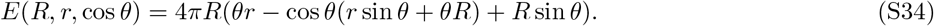

The equilibrium position of the droplet is determined by the *r*_eq_ that minimizes the free energy *E*(*V, r*) for a fixed droplet volume *V*. The constrained optimization problem can be solved by introducing a Lagrange multiplier *λ*:

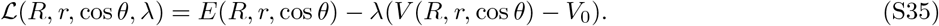

The optimum is given by

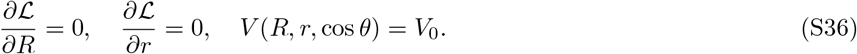

The solutions are:

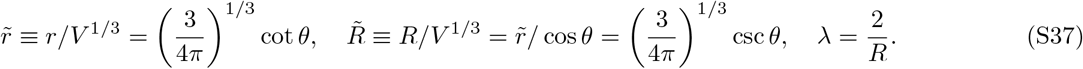

Since the radius *r* must be positive, this solution is only physical for *θ* ∈ (0, *π/*2). For *θ* ∈ (*π/*2, *π*), the surface energy increases monotonically with *r*, and the droplet would prefer to reside at the smallest radius possible. Putting these results together, we arrive at the equilibrium radius of the tubule where the droplet prefers to locate:

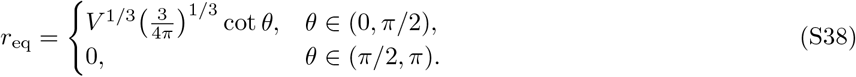

Hence, tuning tether abundance *ψ*_g_ (and thus the contact angle *θ*) can control the equilibrium position of the droplet on a tubule of varying radius (Fig. S1C,D). Theoretical predictions for the *ψ*_g_ values used in Fig.3 in the main text are indicated by red data points: for *ψ*_g_ = 0.05, 0.10, the droplet moves to the left to decrease *r*; for *ψ*_g_ = 0.15, 0.20, 0.25, the droplet moves to the right to increase *r*.

#### B. Droplet migration velocity

As a droplet migrates on a tubule, its velocity 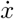is determined by the balance between the driving force due to the surface energy gradient and the drag force:

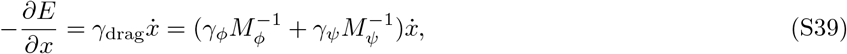

where *γ*_drag_ is the drag coefficient, which we assume to be inversely proportional to the mobility coefficients *M*_*ϕ*_ and *M*_*ψ*_ of the condensate and tethers, respectively. *γ*_*ϕ*_ and *γ*_*ψ*_ are constants that depend on the concentration profile of the condensate and tethers, respectively.

As derived in the previous section *E*(*V, r*) depends on both the droplet volume *V* and the local tubule radius *r*, which gives the driving force 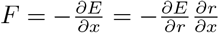. The velocity 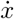depends on *M*_*ψ*_ via an inverse linear relation:

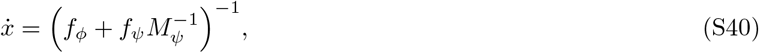

Where

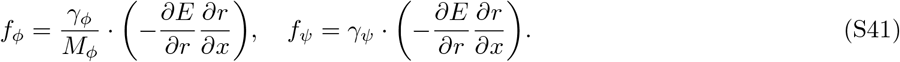

Estimating *f*_*ϕ*_ and *f*_*ψ*_ requires computing the drag coefficients *γ*_*ϕ*_ and *γ*_*ψ*_, which we turn to next. The drag coefficient is determined from the total dissipation:

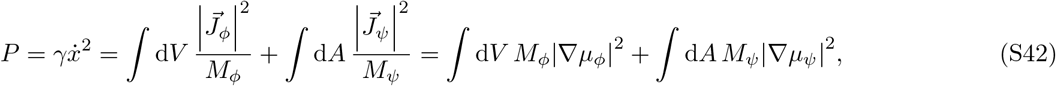

where 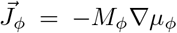 and 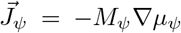are the fluxes of the condensate and tethers, respectively. Further integrating by parts, we have

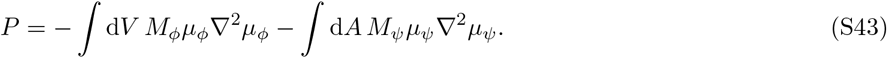

For a traveling wave solution, we have 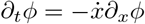and 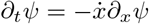. Thus, their (model B) dynamics satisfy

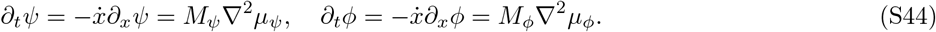

*µ*_*ϕ*,ψ_ are solutions to Poisson equations with source terms 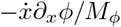 and 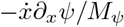, respectively. We can rescale the chemical potentials by the strength of the source:

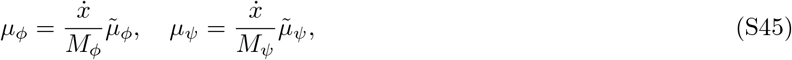

where 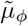 and 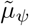 are solutions to the Poisson equations with unit source terms:

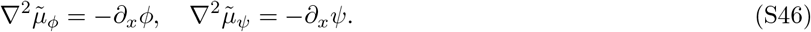

Substituting these back into the expression for *P*, we have

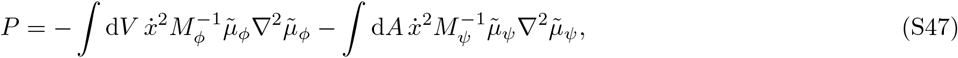

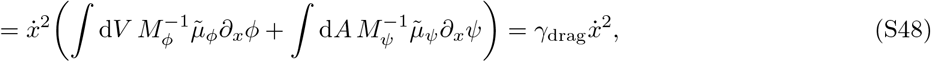

which gives the coefficients

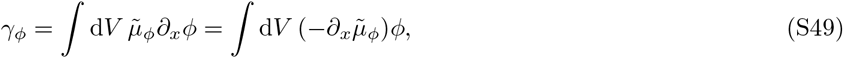

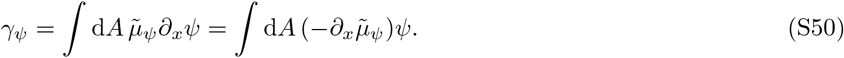

Since the chemical potential gradients 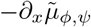 are non-zero only at the interface region, we estimate that *γ*_*ϕ,ψ*_ are proportional to the interface area/length as well as to the concentrations:

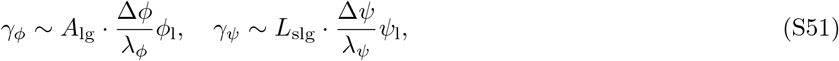

where Δ*ϕ* and Δ*ψ* are the concentration differences across the interface, *λ*_*ϕ*_ and *λ*_*ψ*_ are the interface thicknesses, *A*_lg_ is the area of the condensate-solvent interface, and *L*_slg_ is the length of the three-phase contact line.

For the simulations in Fig. 3E, since the geometry and the condensate concentrations remain approximately unchanged, the main effect of changing tether abundance *ψ*_g_ should be due to the *ψ* dependence in *γ*_*ψ*_. Thus, we estimate:

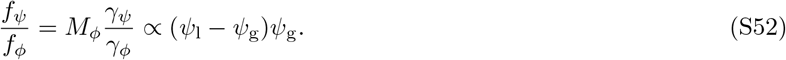

This is consistent with our numerical results (Fig. S2), where we find a linear relation between *f*_*ψ*_*/f*_*ϕ*_ and (*ψ*_l_ − *ψ*_g_)*ψ*_g_.

### IV. DETAILS OF NUMERICAL SIMULATIONS

To solve the equations of motion (Eq. S2) for the condensate and tethers, we use a finite-volume numerical scheme with first-order forward Euler time-stepping for time evolution. The condensate field *ϕ* obeys no-flux boundary conditions at all boundaries in addition to wetting boundary conditions at membrane interfaces. The tether field *ψ* obeys Dirichlet boundary conditions with fixed bulk tether concentration *ψ*_g_. The wetting boundary condition is prescribed using the ghost point method, where the *ϕ* value at each ghost point is interpolated from the two nearest interior points that are not collinear with the ghost point. The volumes near the boundary are treated with the cut-cell method.

For Fig. 1, the simulation was performed in cylindrical coordinates, with spatial discretization 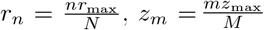, where *r*_max_ = 60 and *z* _max_ = 40 set the system size and *N* = *M* = 128 set the spatial resolution. The system was evolved to its equilibrium state. The contact angle was measured by fitting the contour to a spherical cap *R*^2^ = *r*^2^ + (*z* − *z*_0_)^2^, which gives cos *θ* = −*z*_0_*/R*_0_. The parameters are: *χ*_*ϕ*_ = 2.5, *λ*_*ϕ*_ = 1, *χ*_*ψ*_ = 0, *λ*_*ψ*_ = 0; *h*_1_ = 1 for (B)–(D); *ψ*_g_ = 0.02 and *h*_0_ = 0 for (B); *h*_0_ = −0.2 for (E).

For Fig. 2, the simulation was performed in 2D planar coordinates, with spatial discretization 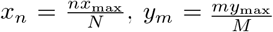 with *x* _max_ = *y*_max_ = 30 and *N* = *M* = 64. The parameters are: *h*_0_ = −0.2, *h*_1_ = 2, *χ*_*ϕ*_ = 2.5, *λ*_*ϕ*_ = 1, *χ*_*ψ*_ = 0, *λ*_*ψ*_ = 0. Tether concentration is fixed at the boundary by Dirichlet boundary condition *ψ*_g_ = 0.05. The videos for the dynamics are available as supplemental videos.

To quantify how fast the droplet reaches its equilibrium configuration at the lower-left corner where the two membranes meet, we defined an average distance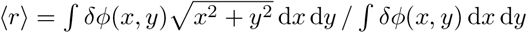, where *δϕ* = *ϕ* − *ϕ*_g_ is the condensate concentration subtracted by the dilute phase.

For Fig. 3 (and Figs. S1 and S2), the simulation was performed in cylindrical coordinates (*r, x*), where the *x*-axis runs along the center of the tubule. The tubule radius is given by *R*(*x*) = *R*_0_ + *R*_1_*x*, with *R*_0_ = 1.5 and *R*_1_ = 1*/*15. Spatial discretization is given by 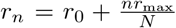, 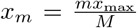, where *r*_0_ = 1.0, *r*_max_ = 15 and *x*_max_ = 30 set the system size and *N* = 64 and *M* = 128 set the spatial resolution. The equations are only evolved in grid points inside the domain [*r > R*(*x*)] with wetting boundary conditions implemented by ghost points. The average position ⟨*x*⟩ is computed from the volume inside the contour *ϕ >* (*ϕ*_l_ + *ϕ*_g_)*/*2. Tether concentration is fixed at the boundary by Dirichlet boundary condition at the right boundary (*x* = *x*_max_). The parameters are: *h*_0_ = − 0.2, *h*_1_ = 1, *χ*_*ϕ*_ = 2.5, *λ*_*ϕ*_ = 1, *χ*_*ψ*_ = 0, *λ*_*ψ*_ = 0. For Fig. 3A, *ψ*_g_ = 0.20 and *M*_*ψ*_ = 1.0; for Fig. 3B, *M*_*ψ*_ = 1.0; for Fig. 3C, *ψ*_g_ = 0.10.

## Footnotes

1 The correction due to this approximation is 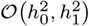, which, as shown below, is a higher-order term.

## References

[1] Y. Shin and C. P. Brangwynne, Liquid phase condensation in cell physiology and disease, Science 357, eaaf4382 (2017).

[2] S. F. Banani, H. O. Lee, A. A. Hyman, and M. K. Rosen, Biomolecular condensates: organizers of cellular biochemistry, Nat. Rev. Mol. Cell Biol. 18, 285 (2017).

[3] X. Su, J. A. Ditlev, E. Hui, W. Xing, S. Banjade, J. Okrut, D. S. King, J. Taunton, M. K. Rosen, and R. D. Vale, Phase separation of signaling molecules promotes t cell receptor signal transduction, Science 352, 595 (2016).

[4] Y. G. Zhao and H. Zhang, Phase separation in membrane biology: The interplay between membrane-bound organelles and membraneless condensates, Dev. Cell 55, 30 (2020).

[5] A. Mangiarotti and R. Dimova, Biomolecular condensates in contact with membranes, Annu. Rev. Biophys. (2024).

[6] L. B. Case, Membranes regulate biomolecular condensates, Nat. Cell Biol. 24, 404 (2022).

[7] H. M. J. Weakly and S. L. Keller, Coupling liquid phases in 3d condensates and 2d membranes: Successes, challenges, and tools, Biophys. J. (2023).

[8] O. Beutel, R. Maraspini, K. Pombo-García, C. Martin-Lemaitre, and A. Honigmann, Phase separation of zonula occludens proteins drives formation of tight junctions, Cell 179, 923 (2019).

[9] S. He, V. L. Crans, and M. C. Jonikas, The pyrenoid: the eukaryotic CO_2_-concentrating organelle, Plant Cell 35, 3236 (2023).

[10] V. Swaminathan, R. S. Fischer, and C. M. Waterman, The FAK-Arp2/3 interaction promotes leading edge advance and haptosensing by coupling nascent adhesions to lamellipodia actin, Mol. Biol. Cell 27, 1085 (2016).

[11] Q. Xiao, C. K. McAtee, and X. Su, Phase separation in immune signalling, Nat. Rev. Immunol. 22, 188 (2022).

[12] J. H. Hennacy, N. Atkinson, A. Kayser-Browne, S. L. Ergun, E. Franklin, L. Wang, S. Eicke, Y. Kazachkova, M. Kafri, F. Fauser, J. Vilarrasa-Blasi, R. E. Jinkerson, S. C. Zeeman, A. J. McCormick, and M. C. Jonikas, Saga1 and mith1 produce matrix-traversing membranes in the CO_2_-fixing pyrenoid, Nat. Plants 10, 2038 (2024).

[13] M. T. Meyer, A. K. Itakura, W. Patena, L. Wang, S. He, T. Emrich-Mills, C. S. Lau, G. Yates, L. C. M. Mackinder, and M. C. Jonikas, Assembly of the algal CO_2_-fixing organelle, the pyrenoid, is guided by a rubisco-binding motif, Sci. Adv. 6, eabd2408 (2020).

[14] T. Young, Iii. an essay on the cohesion of fluids, Philosophical transactions of the royal society of London, 65 (1805).

[15] B. D. Cassie and S. Baxter, Wettability of porous surfaces, Trans. Faraday Soc. 40, 546 (1944).

[16] B. Andreotti and J. H. Snoeijer, Statics and dynamics of soft wetting, Annu. Rev. Fluid Mech. 52, 285 (2020).

[17] P. C. Hohenberg and B. I. Halperin, Theory of dynamic critical phenomena, Rev. Mod. Phys. 49, 435 (1977).

[18] S. Mao, D. Kuldinow, M. P. Haataja, and A. Košmrlj, Phase behavior and morphology of multicomponent liquid mixtures, Soft Matter 15, 1297 (2019).

[19] H. Hofmann, A. Soranno, A. Borgia, K. Gast, D. Nettels, and B. Schuler, Polymer scaling laws of unfolded and intrinsically disordered proteins quantified with singlemolecule spectroscopy, Proc. Natl. Acad. Sci. U. S. A. 109, 16155 (2012).

[20] H. Wang, F. M. Kelley, D. Milovanovic, B. S. Schuster, and Z. Shi, Surface tension and viscosity of protein condensates quantified by micropipette aspiration, Biophysical Reports 1 (2021).

[21] D. N. Itzhak, S. Tyanova, J. Cox, and G. H. Borner, Global, quantitative and dynamic mapping of protein subcellular localization, eLife 5, e16950 (2016).

[22] E. S. Freeman Rosenzweig, B. Xu, L. Kuhn Cuellar, Martinez-Sanchez, M. Schaffer, M. Strauss, H. N. Cartw right, P. Ronceray, J. M. Plitzko, F. Förster, N. S. Wingreen, B. D. Engel, L. C. M. Mackinder, and M. C. Jonikas, The eukaryotic CO_2_-concentrating organelle is liquid-like and exhibits dynamic reorganization, Cell 171, 148 (2017).

[23] B. J. Carroll, The accurate measurement of contact angle, phase contact areas, drop volume, and laplace excess pressure in drop-on-fiber systems, J. Colloid Interface Sci. 57, 488 (1976).

[24] G. Mchale, M. I. Newton, and B. J. Carroll, The shape and stability of small liquid drops on fibers, Oil Gas Sci. Technol. 56, 47 (2001).

[25] J. Van Hulle, F. Weyer, S. Dorbolo, and N. Vandewalle, Capillary transport from barrel to clamshell droplets on conical fibers, Phys. Rev. Fluids 6, 024501 (2021).

[26] H. B. Eral, J. de Ruiter, R. de Ruiter, J. M. Oh, C. Semprebon, M. Brinkmann, and F. Mugele, Drops on functional fibers: from barrels to clamshells and back, Soft Matter 7, 5138 (2011).

[27] T. GrandPre, Y. Zhang, A. G. T. Pyo, B. Weiner, J.-L. Li, M. C. Jonikas, and N. S. Wingreen, Impact of linker length on biomolecular condensate formation, PRX Life 1, 023013 (2023).

[28] T. Litschel, C. F. Kelley, X. Cheng, L. Babl, N. Mizuno, L. B. Case, and P. Schwille, Membrane-induced 2d phase separation of the focal adhesion protein talin, Nat. Commun. 15, 4986 (2024).

[29] L. B. Case, M. De Pasquale, L. Henry, and M. K. Rosen, Synergistic phase separation of two pathways promotes integrin clustering and nascent adhesion formation, eLife 11, e72588 (2022).

[30] S. Sun, T. GrandPre, D. T. Limmer, and J. T. Groves, Kinetic frustration by limited bond availability controls the lat protein condensation phase transition on membranes, Sci. Adv. 8, eabo5295 (2022).

[31] L. B. Case, X. Zhang, J. A. Ditlev, and M. K. Rosen, Stoichiometry controls activity of phase-separated clusters of actin signaling proteins, Science 363, 1093 (2019).

[32] Z. C. Scott, K. Koning, M. Vanderwerp, L. Cohen, L. M. Westrate, and E. F. Koslover, Endoplasmic reticulum network heterogeneity guides diffusive transport and kinetics, Biophys. J. 122, 3191 (2023).

[33] K. Speckner, L. Stadler, and M. Weiss, Unscrambling exit site patterns on the endoplasmic reticulum as a quenched demixing process, Biophys. J. 120, 2532 (2021).

[34] W. van Leeuwen, D. T. M. Nguyen, R. Grond, T. Veenendaal, C. Rabouille, and G. G. Farías, Stress-induced phase separation of eres components into sec bodies precedes er exit inhibition in mammalian cells, J. Cell Sci. 135, jcs260294 (2022).

[35] C. E. Cornell, A. D. Skinkle, S. He, I. Levental, K. R. Levental, and S. L. Keller, Tuning length scales of small domains in cell-derived membranes and synthetic model membranes, Biophys. J. 115, 690 (2018).

[36] Q. Yu and A. Košmrlj, Pattern formation of lipid domains in bilayer membranes, Soft Matter 21, 4288 (2025).

[37] T.-S. Hsieh, V. A. Lopez, M. H. Black, A. Osinski, K. Pawlowski, D. R. Tomchick, J. Liou, and V. S. Tagliabracci, Dynamic remodeling of host membranes by self-organizing bacterial effectors, Science 372, 935 (2021).

[38] J. Tjalma, V. Galstyan, J. Goedhart, L. Slim, N. B. Becker, and P. R. Ten Wolde, Trade-offs between cost and information in cellular prediction, Proc. Natl. Acad. Sci. U. S. A. 120, e2303078120 (2023).

[39] D. Hathcock, Q. Yu, and Y. Tu, Time-reversal symmetry breaking in the chemosensory array reveals a general mechanism for dissipation-enhanced cooperative sensing, Nat. Commun. 15, 8892 (2024).

## REFERENCES

[1] S. Mao, D. Kuldinow, M. P. Haataja, and A. Košmrlj, Phase behavior and morphology of multicomponent liquid mixtures, Soft Matter 15, 1297 (2019).

[2] P. C. Hohenberg and B. I. Halperin, Theory of dynamic critical phenomena, Rev. Mod. Phys. 49, 435 (1977).

[3] H. Hofmann, A. Soranno, A. Borgia, K. Gast, D. Nettels, and B. Schuler, Polymer scaling laws of unfolded and intrinsically disordered proteins quantified with single-molecule spectroscopy, Proc. Natl. Acad. Sci. U. S. A. 109, 16155 (2012).

[4] H. Wang, F. M. Kelley, D. Milovanovic, B. S. Schuster, and Z. Shi, Surface tension and viscosity of protein condensates quantified by micropipette aspiration, Biophysical Reports 1 (2021).

[5] D. N. Itzhak, S. Tyanova, J. Cox, and G. H. Borner, Global, quantitative and dynamic mapping of protein subcellular localization, eLife 5, e16950 (2016).

[6] B. J. Carroll, The accurate measurement of contact angle, phase contact areas, drop volume, and laplace excess pressure in drop-on-fiber systems, J. Colloid Interface Sci. 57, 488 (1976).

